# Comparative analysis of N-terminal cysteine dioxygenation and prolyl-hydroxylation as oxygen sensing pathways in mammalian cells

**DOI:** 10.1101/2023.06.25.545688

**Authors:** Ya-Min Tian, Philip Holdship, Trang Quynh To, Peter J Ratcliffe, Thomas P Keeley

**Affiliations:** Target Discovery Institute, Nuffield Department of Medicine, University of Oxford, Oxford OX3 7FZ, UK; Ludwig Institute for Cancer Research, Nuffield Department of Medicine, University of Oxford, Oxford OX3 7FZ, UK; Department of Earth Sciences, University of Oxford, South Parks Rd, Oxford, OX1 3AN, UK; The Francis Crick Institute, 1 Midland Road, London NW1 1AT, UK

**Keywords:** hypoxia, sensing, dioxygenase, hydroxylase, ADO, HIF, PHD

## Abstract

In animals, adaptation to changes in cellular oxygen levels is coordinated largely by the 2-oxoglutarate dependent prolyl-hydroxylase domain (PHD) dioxygenase family, which regulate the stability of their hypoxia-inducible factor (HIF) substrates to promote expression of genes that adapt cells to hypoxia. Recently, 2-aminoethanethiol dioxygenase (ADO) was identified as a novel O_2_-sensing enzyme in animals. Through N-terminal cysteine dioxygenation and the N-degron pathway, ADO regulates the stability of a set of non-transcription factor substrates; the regulators of G-protein signalling 4, 5 and 16, and interleukin-32. Here, we set out to compare and contrast the *in cellulo* characteristics of ADO and PHD enzymes in an attempt to better understand their co-evolution in animals. We find that ADO operates to regulate the stability of its substrates rapidly and with similar O_2_-sensitivity to the PHD/HIF pathway. ADO appeared less sensitive to iron chelating agents or transition metal exposure than the PHD enzymes, possibly due to tighter catalytic-site Fe^2+^ coordination. Unlike the PHD/HIF pathway, the ADO/N-degron pathway was not subject to feedback by hypoxic induction of ADO and induction of ADO substrates was well sustained in response to prolonged hypoxia. The data also reveal strong interactions between proteolytic regulation of targets by ADO and transcriptional induction of those targets, that shape integrated cellular responses to hypoxia.

## Introduction

Systems that sense, adapt and alleviate exposure to hypoxia (low O_2_) are observed throughout evolution, reflecting the fundamental importance of O_2_ in biology and its obligate role in multicellular life forms. In mammals, a sensing system comprised of set of 2-oxoglutarate dependent dioxygenase enzymes (<u>P</u>rolyl <u>H</u>ydroxylase <u>D</u>omain, PHD1, 2, and 3, otherwise known as EGLN 2,1, 3) and their Hypoxia-Inducible factor (HIF-1, -2, and 3α) target substrates is widespread and likely responsible for the majority of adaptive transcriptional responses to hypoxia (reviewed in (1,2)). Similarly, oxygen sensing in plants is orchestrated by plant cysteine dioxygenases (PCOs) which regulate the stability of ethylene response factor VII transcription factors (3-5). Both systems respond to changes in environmental pO_2_ by inducing a pattern of gene expression capable of reducing cellular O_2_ usage, whilst simultaneously increasing O_2_ delivery at the tissue level.

It was recently demonstrated that enzymatic cysteine dioxygenation also occurs in mammals through an orthologue of PCO, 2-aminoethanethiol dioxygenase (ADO) (6). Together, these enzymes constitute the N-terminal cysteine dioxygenase family and regulate protein stability via the Arg/Cys branch of the N-degron pathway (7). Unlike PCO and PHD enzymes, the known substrates of ADO are not transcription factors but rather regulators of G-protein signalling (RGS4, 5 and 16) and the atypical cytokine interleukin-32 (IL-32). These RGS proteins interact with Gα*q* and *i* subunits of the heterotrimeric G-protein complex to increase the rate of GTP hydrolysis, thereby repressing downstream signalling through the associated G-protein coupled receptor (as reviewed in (8)). The physiological role of IL-32 remains to be elucidated, but strong associations have been made between its expression and the severity of non-alcoholic fatty liver disease (9,10). IL-32 has also been implicated in the host response to hepatitis B and C infections (11,12). Under normoxic conditions, ADO substrates are degraded via N-degron mediated proteolysis, but reductions in cellular O_2_ levels lead to reduction in ADO activity and thus to substrate stabilisation. Much like the PHD enzymes, recombinant ADO and PCO are highly sensitive to O_2_ *in vitro* (6,13) and so could potentially transduce signals under physiological and pathophysiological levels of hypoxia (e.g. 0-10% O_2_). However, in contrast with the detailed characterisation of PHD/HIF little is known about the analogous ADO/RGS system, or how the two systems interface.

Here we compare and contrast the operation of these O_2_ sensing pathways in mammalian cells. We show that both pathways operate with similar *in cellulo* sensitivity with respect to O_2_, but have important biochemical and pharmacological distinctions. Understanding the constraints within which each pathway operates in cells will be important in deciphering their shared and distinct roles in mammalian hypoxia physiology and potentially in the therapeutic modulation of these systems.

## Results

A defining feature of physiological O_2_ sensing mechanisms, exemplified by the HIF/PHD system, is a graded response to changes in cellular oxygenation within the physiological range. In *in vitro* studies, both the PHD enzymes and ADO have been reported to be similarly sensitive to O_2_, with apparent K_m_O_2_ values 200-500μM O_2_ (6,14,15). We have previously reported limited studies of the sensitivity of the ADO substrates, RGS4 and 5 to hypoxia in cell lines. In the present analysis we first sought to extend this work by comparing the oxygen sensitivity of known ADO substrates to that of HIF over a wide range of physiological to pathophysiological oxygen concentrations. Since cellular oxygenation in culture is defined by a complex interaction between the ambient O_2_ levels, monolayer confluency and cellular O_2_ consumption (16), these experiments were conducted under similar conditions across a range of cell lines (SH-SY5Y, RKO, HepG2, Kelly and EA.hy926). To guide these analyses, RNA extracts from each cell line were initially screened for the presence of transcripts encoding the 4 known ADO substrates; *RGS4, RGS5, RGS16* and *IL32* (Fig. 1A). This revealed that, amongst the cell lines studied, RGS4 is the most widely expressed substrate, with *RGS16* and *IL32* being confined to the neuroblastomal (SH-SY5Y and Kelly) and the gastrointestinal cell lines (HepG2 and RKO), respectively. We then exposed the cells to different O_2_ levels (18, 7.5, 3, 1 and 0.1% O_2_) for 4 hours (for RGS4/5) or 18 hours (for IL-32, for time course of induction see Figure 6) and measured the abundance of HIF-1α and expressed ADO substrates. These experiments revealed strikingly similar sensitivity of HIF-1α and RGS4/5 and/or IL-32 to hypoxia in all cell types studied. Data is illustrated for all proteins studied in each cell line in Figure 1B and quantified using non-linear regression analysis in Figure 1C, and demonstrates a closely similar relationship between HIF-1α and ADO substrate protein levels and applied pO_2_. We were unable to detect RGS16 protein in Kelly cells, nor was RGS5 protein detectable in EA.hy926 cells, despite the presence of the corresponding mRNA transcript in these cells.

**Figure 1.**
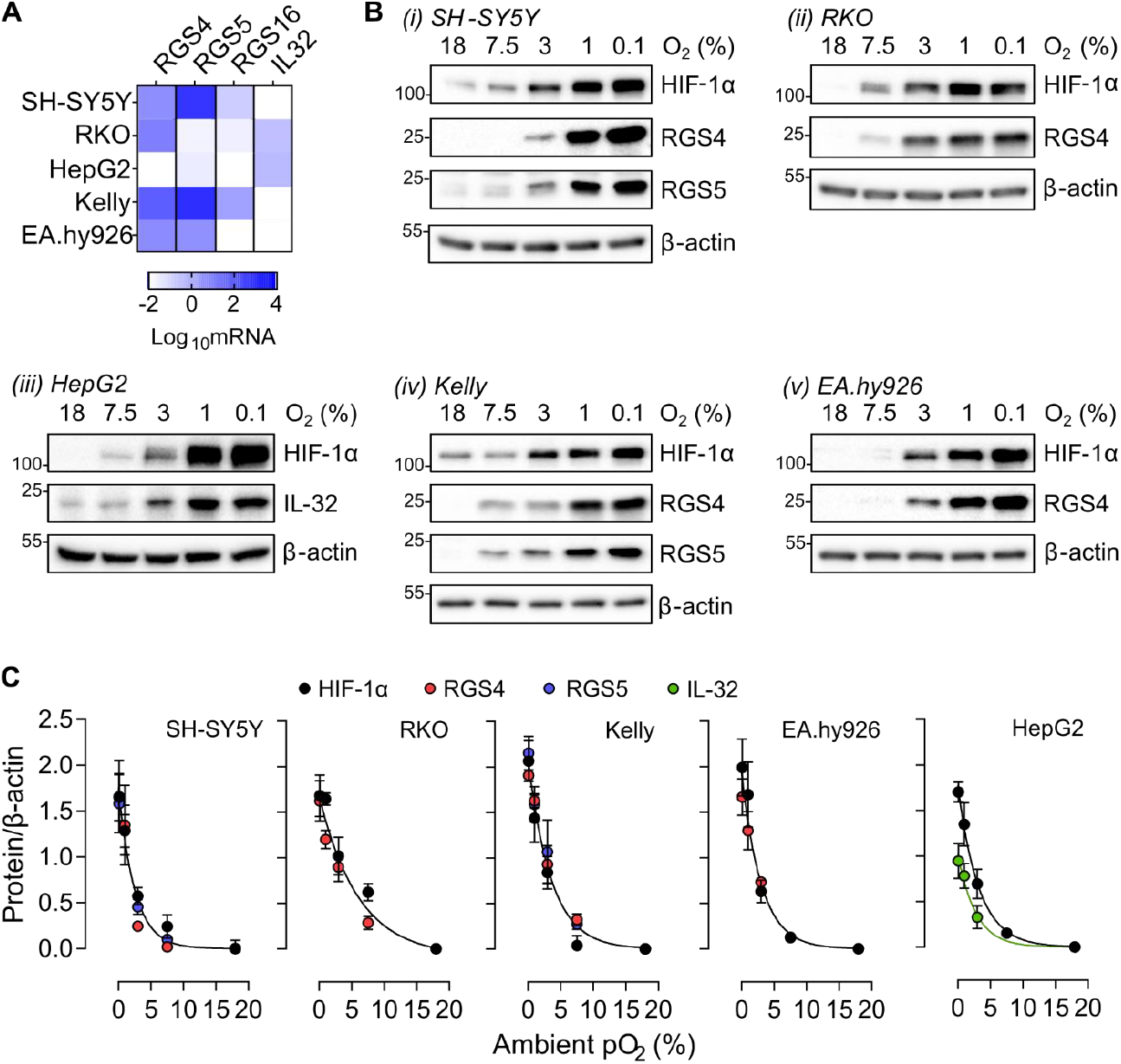
Response to graded hypoxia in mammalian cell lines. (A) The expression pattern of ADO substrate mRNA transcripts in 5 cell lines. Colour scale (white to blue) represents the Log10 fold change in substrate mRNA level relative to HPRT. (B) SH-SY5Y, RKO, HepG2, Kelly and EA.hy926 cells were subjected to 4h hypoxia (18h for IL-32 in HepG2) at the indicated level of hypoxia and expression of ADO substrates present in each cell line assessed by immunoblotting. HIF-1α protein levels were analysed in parallel for comparison. (C) Immunoblots from B were quantified and subjected to non-linear regression analysis. In all but HepG2 cells, all data points were better represented statistically using one curve than individual curves, with P values 0.15-0.86. Data represent the mean ± S.D. from 3 independent experiments.

In previous studies, the destabilizing action of N-terminal cysteine dioxygenation in human cells was demonstrated by measuring the half-life of fusion proteins bearing the N-terminus of the plant ERF-VII transcription factor fused to a green fluorescent protein-V5 reporter protein (6). HIF-1α has been reported as having a remarkably short half-life in oxygenated cells (17). Given similarities between HIF-1α, RGS4 and 5 with respect to protein accumulation in hypoxic cells, we next wanted to compare the relative level of these proteins in cells using assays of endogenous protein levels following re-oxygenation of hypoxic cells (Fig. 2). Whilst levels of RGS4 and RGS5 proteins remained stable under continued hypoxic exposure, reoxygenation resulted in rapid degradation of RGS4 and 5 with a T_1/2_ (time for disappearance of half of the species) of approximately 4 minutes. These kinetics were very similar to those of HIF-1α (T_1/2_ 5 minutes), assayed under the same conditions. Notably, these experiments were performed in the absence of the protein synthesis inhibitor, cycloheximide, as this was found to reduce RGS4 and 5 expression beyond reliable detection at all time points. Therefore, the figures for T_1/2_ represent maximum values.

**Figure 2.**
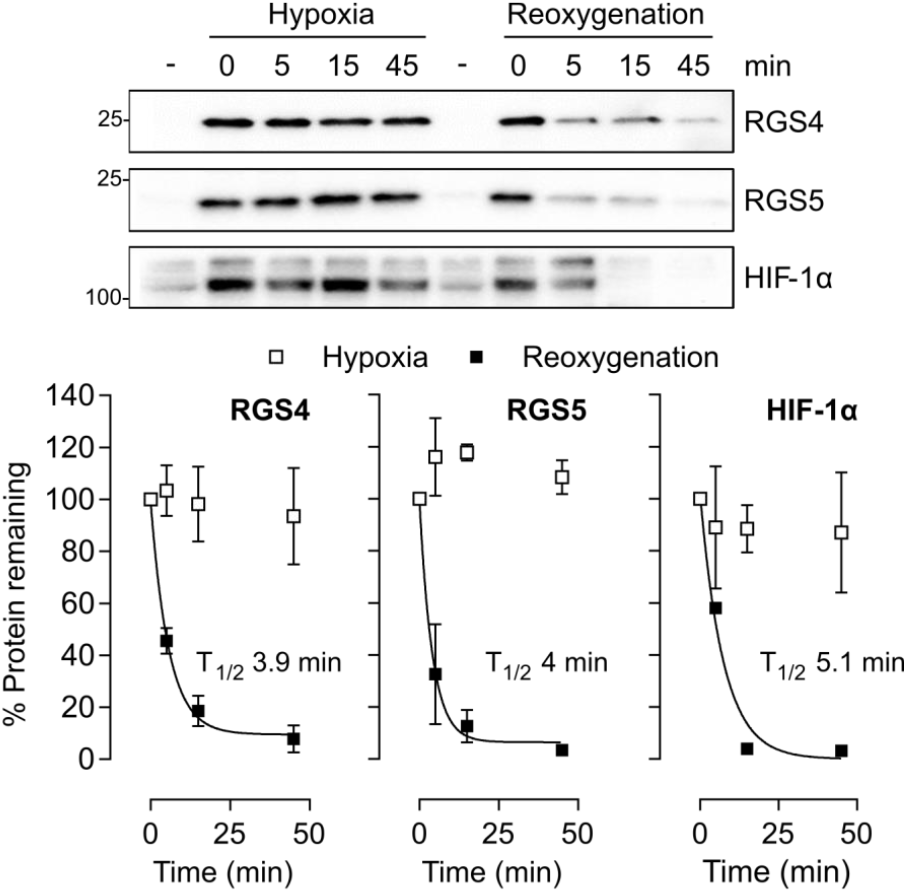
Degradation of RGS4, 5 and HIF-1α in the presence of oxygen. SH-SY5Y cells were cultured under hypoxic conditions (1% O2) for 16h then either maintained under hypoxia or reoxygenated by exposure to atmospheric levels of oxygen for 5, 15 or 45min. Levels of RGS4, 5 and HIF-1α protein were assessed by immunoblotting, quantified and subjected to non-linear exponential decay regression to quantify the degradation. Data plotted are mean ± S.D. from 3 independent experiments.

The HIF Prolyl-hydroxylases and Nt-Cysteine dioxygenases share a similar His-Asp-His motif for Fe^2+^ coordination (18,19). It was therefore surprising that in previous studies ADO appeared sensitive to a narrower range of Fe^2+^ chelating small molecules than the PHDs (6). To investigate this further, we tested a range of iron chelating compounds for their ability to inhibit ADO using RGS4_1-11_GFP fusion protein that has been shown to accurately report ADO activity in cells (20). RKO cells stably expressing RGS4_1-11_GFP were treated with 6 different iron chelators (2,2DIP -2,2′-dipyridyl, DFO – deferoxamine, Dp44mT - di-2-pyridylketone 4,4-dimethyl-3-thiosemicarbazone, ICL670A - desferasirox, L1 – deferiprone, SIH - salicylaldehyde isonicotinoyl hydrazone) at a range of concentrations, and accumulation of the reporter assayed by monitoring GFP fluorescence. Of the compounds tested only 2,2DIP, Dp44mT and SIH manifest inhibition of ADO at concentrations that did not elicit cytotoxicity (Fig. 3A). Similar results were obtained when the accumulation of endogenous RGS4 or 5 was analysed in RKO or SH-SY5Y cells, respectively (Fig. 3B). In contrast, all compounds were broadly capable of stabilising HIF-1α (Fig. 3B). Analysis of intracellular free Fe^2+^ in RKO cells using a fluorescent indicator demonstrated that all compounds except L1 reduced intracellular Fe^2+^ levels within 4 hours of treatment (Fig. 3C). Whilst this was consistent with the presence or absence of HIF-1α stabilisation in this cell type (Fig. 3B), efficacy of ADO inhibition (as inferred from RGS4 expression) did not correlate with intracellular free Fe^2+^ chelation, suggesting a more complex interface between chelators and the catalytic iron centre. To investigate this further, we next analysed the total Fe content of FLAG-tagged ADO and PHD2 proteins immunoprecipitated from SH-SY5Y cells treated with either 2,2DIP or DFO. In line with observations on RGS4/5 stability, 2,2DIP was potent at removing iron from both ADO and PHD2, whereas DFO had very little impact on ADO iron content despite effectively chelating all the iron from PHD2 (Fig. 3D). Interestingly, ADO consistently contained a higher relative iron content than PHD2 following immunoprecipitation, corroborating earlier observations noting a remarkably high iron occupancy of purified ADO relative to CDO1 (21). Finally, there was no further stabilisation of RGS4 observed in ADO deficient cells following treatment with 2,2DIP, Dp44mT or SIH (Fig. 3E), consistent with an action through ADO.

**Figure 3.**
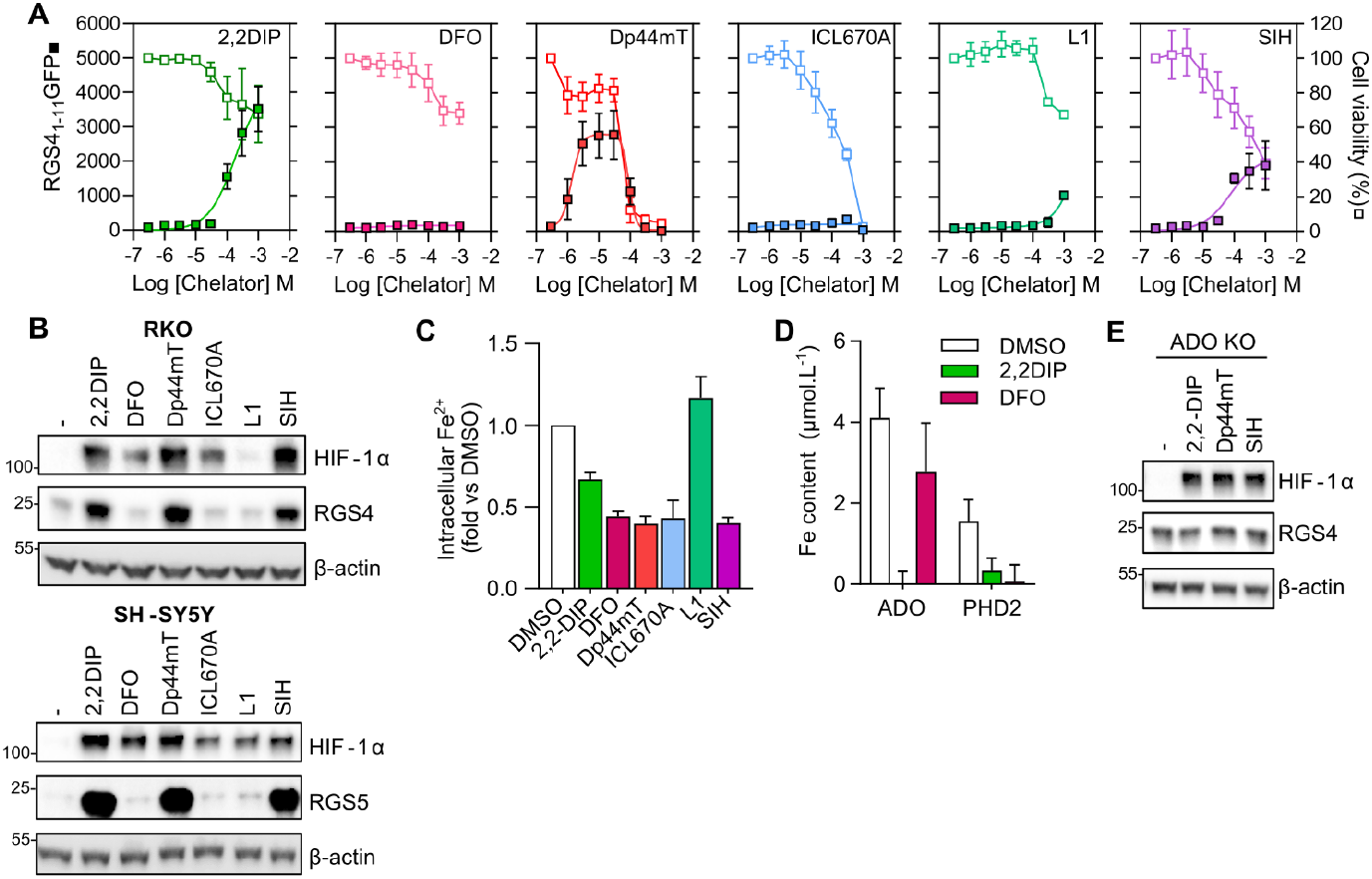
Differential sensitivity of ADO and PHD enzymes to iron chelators. (**A**) RKO cells stably expressing an RGS4_1-11_GFP fusion reporter protein were treated with various iron chelator compounds at the concentrations indicated for 24h. Fluorescence (expressed as arbitrary units, closed squares) was measured and then cellular viability (open squares) assessed using an MTT (3-(4,5-dimethylthiazol-2-yl)-2,5-diphenyltetrazolium bromide) assay. (**B**) RKO and SH-SY5Y cells were treated with the same 6 iron chelating compounds for 4h, at a maximal non-toxic concentration (all 100μM except Dp44mT - 3μM and ICL670A - 30μM), and levels of the endogenous substrates RGS4 or RGS5 assessed by immunoblotting. Expression of HIF-1α was assayed in parallel for comparison. (**C**) Intracellular Fe^2+^ levels were measured using FerroOrange in RKO cells treated with the indicated iron chelators for 4h. (**D**) Measurement of total iron content in FLAG-ADO and FLAG-PHD2 immunoprecipitates from cells treated with DFO or 2,2DIP for 4h, determined using ICP-MS. (**E**) ADO-deficient RKO cells were treated with the Fe^2+^ chelators shown to stabilise RGS4 (2,2DIP, Dp44mT and SIH) for 4h and levels of RGS4 protein assayed. Data in A, C and D represent the mean ± S.D. from 3 independent experiments; all immunoblots are representative of at least 3 independent experiments.

In addition to Fe^2+^, the PHDs and other 2-OG dependent dioxygenases are inhibited by transition metal ions such as Co^2+^ (22-24) and activation of HIF target genes such as erythropoietin is an important feature of the medical toxicology of such metals. The proposed explanation for the inhibition of PHD enzymes by Co^2+^ is displacement of Fe^2+^ within the catalytic domain, thus rendering the enzyme inactive (24). In view of the differences in the efficacy of Fe^2+^ chelators observed earlier, we hypothesised that ADO might be differentially sensitive to Co^2+^, or other divalent cations, as compared to the PHD enzymes. Treatment of RGS4_1-11_GFP reporter cells with increasing concentrations of either CoCl_2_, MnCl_2_ or NiCl_2_ resulted in dose-dependent cytotoxicity, with minimal accumulation of GFP (Fig. 4A). However, CoCl_2_ treatment did elicit striking induction of endogenous RGS4 in RKO cells and RGS5 in SH-SY5Y cells (Fig. 4B). Note, RGS4 protein level was not assessed in SH-SY5Y cells due to the contending influence of Co^2+^ on HIF-mediated transcription in this cell line. Surprisingly, another ADO target, IL-32, was unaffected and induction of RGS4/5 was also clearly evident in ADO-deficient cells. Induction of RGS4 was not accompanied by detectable changes in *RGS4* mRNA levels, as might be expected if it reflected a transcriptional response to HIF (Fig. 4C).

**Figure 4.**
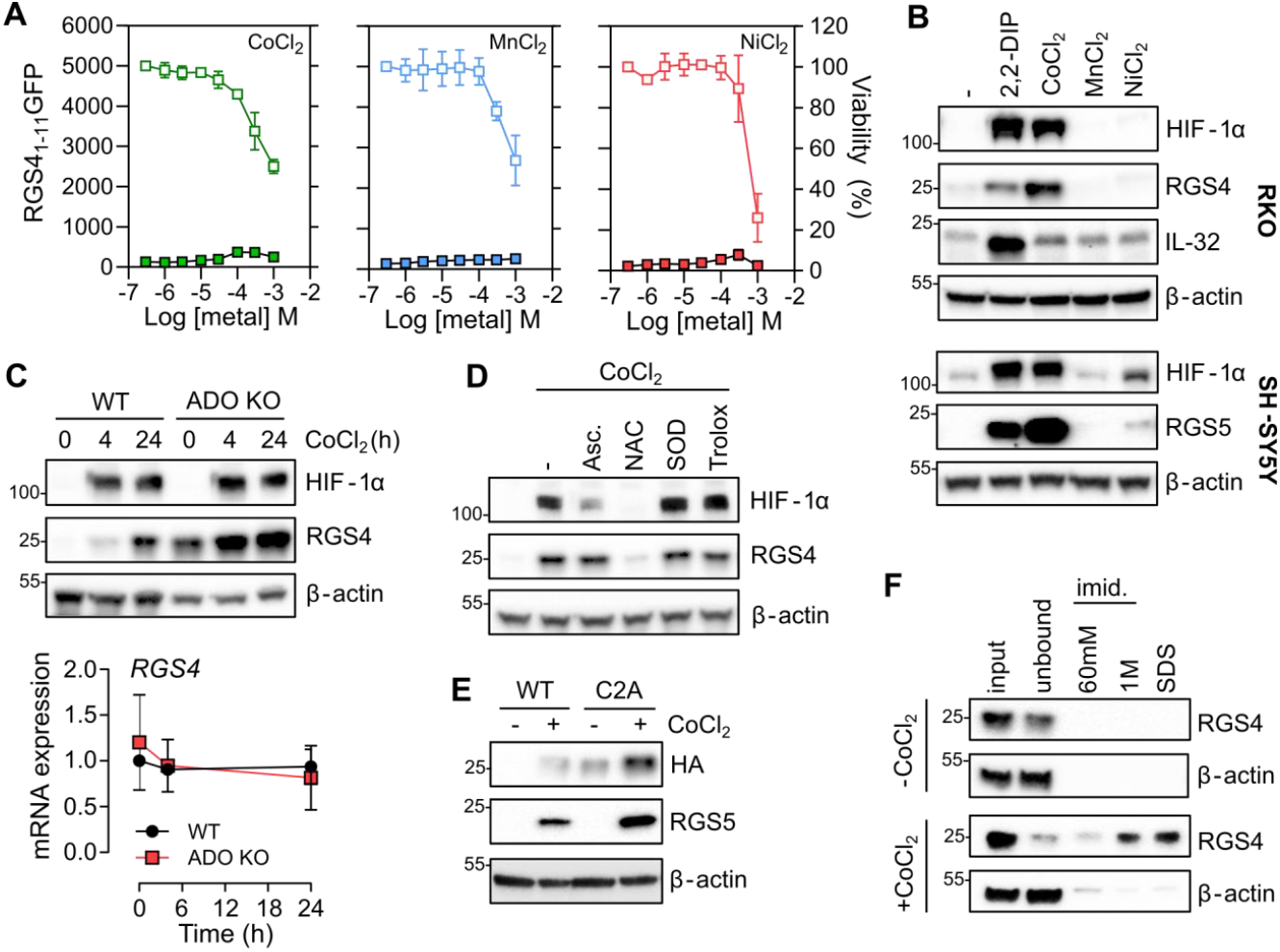
Regulation of RGS4 and 5 by Co^2+^. (**A**) RKO cells stably expressing an RGS4_1-11_GFP fusion reporter gene were treated with increasing concentrations of CoCl_2_, MnCl_2_ or NiCl_2_ for 24h. Accumulation of reporter gene was monitored by fluorescence, and viability assessed in parallel. (**B**) Co^2+^ but not Mn^2+^ or Ni^2+^ induced RGS4 or 5 in RKO and SH-SH5Y cells, respectively. 2,2DIP and HIF-1α immunoblots were used as positive controls. All chemicals were applied at 100μM. (**C**) ADO-competent or -deficient RKO cells were treated with CoCl_2_ for 4 or 24h and accumulation of RGS4 protein or mRNA transcript assessed. (**D**) RKO cells were treated for 24h with Co^2+^ alone or in the presence of ascorbate (asc, 100μM), N-acetyl-L-cysteine (NAC, 1mM), PEGylated superoxide dismutase (SOD, 50U/ml) or the vitamin E analogue Trolox (200μM). (**E**) SH-SY5Y cells stably expressing a C-terminal HA-tagged WT or C2A mutant RGS4 were treated with CoCl_2_ for 24h as before. Endogenous RGS5 levels are shown for comparison. (**F**) RGS4 precipitation from ADO-deficient SH-SY5Y cell extracts using His-beads pre-charged with CoCl_2_ and successive elutions of increasing strength to assess affinity of binding. β-actin was used as a negative control. All data are representative of at least 3 independent experiments.

It was recently proposed that Co^2+^ can cause ‘over-oxidation‘ of Nt-Cysteine residues, targeting them for lysosomal degradation (25). As was reported (25), we also observed an inhibitory effect of N-acetyl cysteine (NAC) on Co^2+^ stimulated RGS4 and also on HIF-1α induction (Fig. 4D). However other compounds known to scavenge reactive oxygen species (ROS) such as ascorbate, the vitamin E analogue Trolox (6-hydroxy-2,5,7,8-tetramethylchroman-2-carboxylic acid), or PEGylated superoxide dismutase had no effect on RGS4 levels. In contrast ascorbate did substantially reduce the level of HIF-1α, consistent with its known function in maintaining activity of the catalytic Fe^2+^ centre of PHD enzymes (26). A persistent action of Co^2+^ in the absence of ADO, in combination with the lack of effect on accumulation of the RGS4_1-11_GFP reporter protein, suggested that Co^2+^ does not interfere with N-terminal cysteine dioxygenation. In line with this hypothesis, Co^2+^ induced protein levels of both RGS4:HA and a mutant that ablated the target cysteine residue RGS4(C2A):HA proteins in SH-SY5Y cells (Fig. 4E), with the latter being more abundant at baseline, as expected from a lack of action of ADO (6). It was previously reported that Co^2+^ could directly bind to HIFα polypeptides and inhibit degradation irrespective of any effect on the PHD enzymes or HIF prolyl hydroxylation (27,28). Interestingly, we found significant Co^2+^-dependent enrichment of RGS4 in His-immunoprecipates from ADO-deficient SH-SY5Y lysates (Fig. 4F), strongly suggesting that RGS4 protein also binds Co^2+^. Taken together, this data indicates a hitherto undescribed effect of Co^2+^ on RGS4 and RGS5 that is apparently independent of ADO and the action of reactive oxygen species.

Another characteristic of the HIF/PHD system is the operation of a feedback loop by which HIF-dependent upregulation of the HIF prolyl hydroxylases PHD2 and PHD3 serves to limit HIF activity over a period of hours (29). We observed that neither ADO protein nor mRNA levels were affected by periods of up to 48 hours of hypoxia in SH-SY5Y cells, directly contrasting with a robust upregulation of PHD3 under those conditions (Fig. 5A and B). Moreover, we found no evidence that other known components of the Arg/Cys N-degron pathway were altered at the mRNA level by hypoxia (Fig. 5C). To address the effects on the time-dependent accumulation of ADO substrate proteins, cell lines were subjected to hypoxia (1% O_2_), over periods ranging from 2 to 48 hours and the levels of substrate mRNAs and proteins assessed (Fig. 6). These experiments revealed rapid accumulation of RGS4 and 5 in all cell lines tested, which was well-sustained even after 48 hours continuous exposure to hypoxia, in contrast with HIF-1α. Most of these responses to hypoxia occurred without change in the corresponding mRNA, consistent with an action of ADO on protein stability. In some cases, a modest induction of the corresponding mRNA was also observed (Fig. 6A, B and D) though this was nearly always delayed relative to the accumulation of protein.

**Figure 5.**
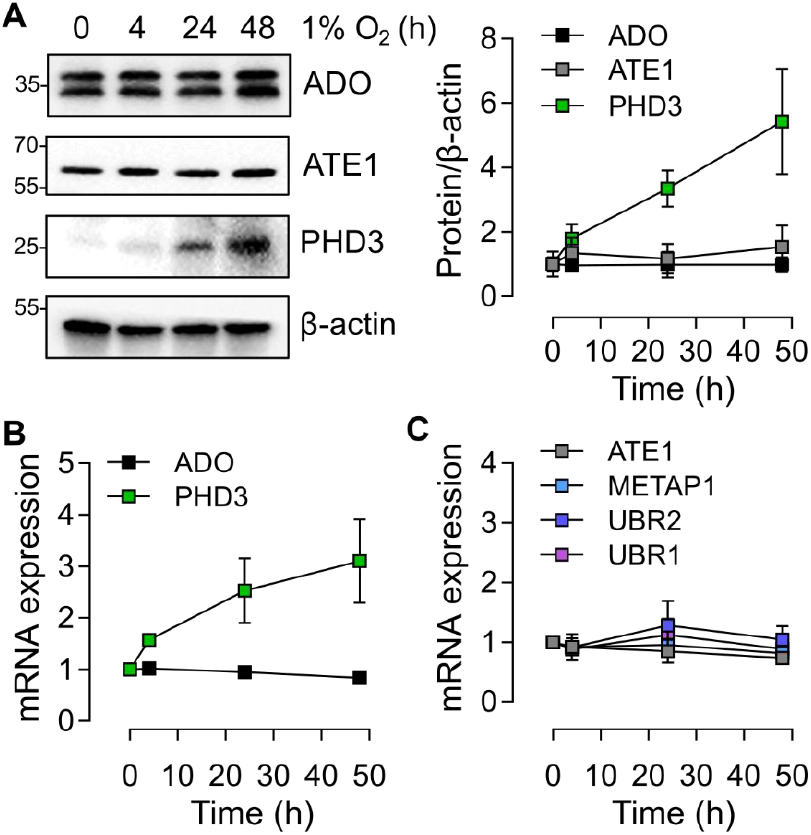
Lack of apparent feedback regulation in the ADO pathway during hypoxic exposure. (**A**) Levels of ADO, arginyl transferase 1 (ATE1) and HIF prolyl-hydroxylase 3 (PHD3) protein in SH-SY5Y exposed to hypoxia for 4, 24 or 48h. (**B**) Hypoxic upregulation of mRNA transcript encoding *PHD3*, but not *ADO*, in SH-SY5Y cells. (**C**) No effect of hypoxia was observed on the transcript levels of other components of the Arg/Cys N-degron pathway (METAP1, methionine aminotransferase 1, UBR1/2, Ubiquitin Protein Ligase E3 Component N-Recognin 1/2). All data represent the mean ± S.D. from 3 independent experiments.

**Figure 6.**
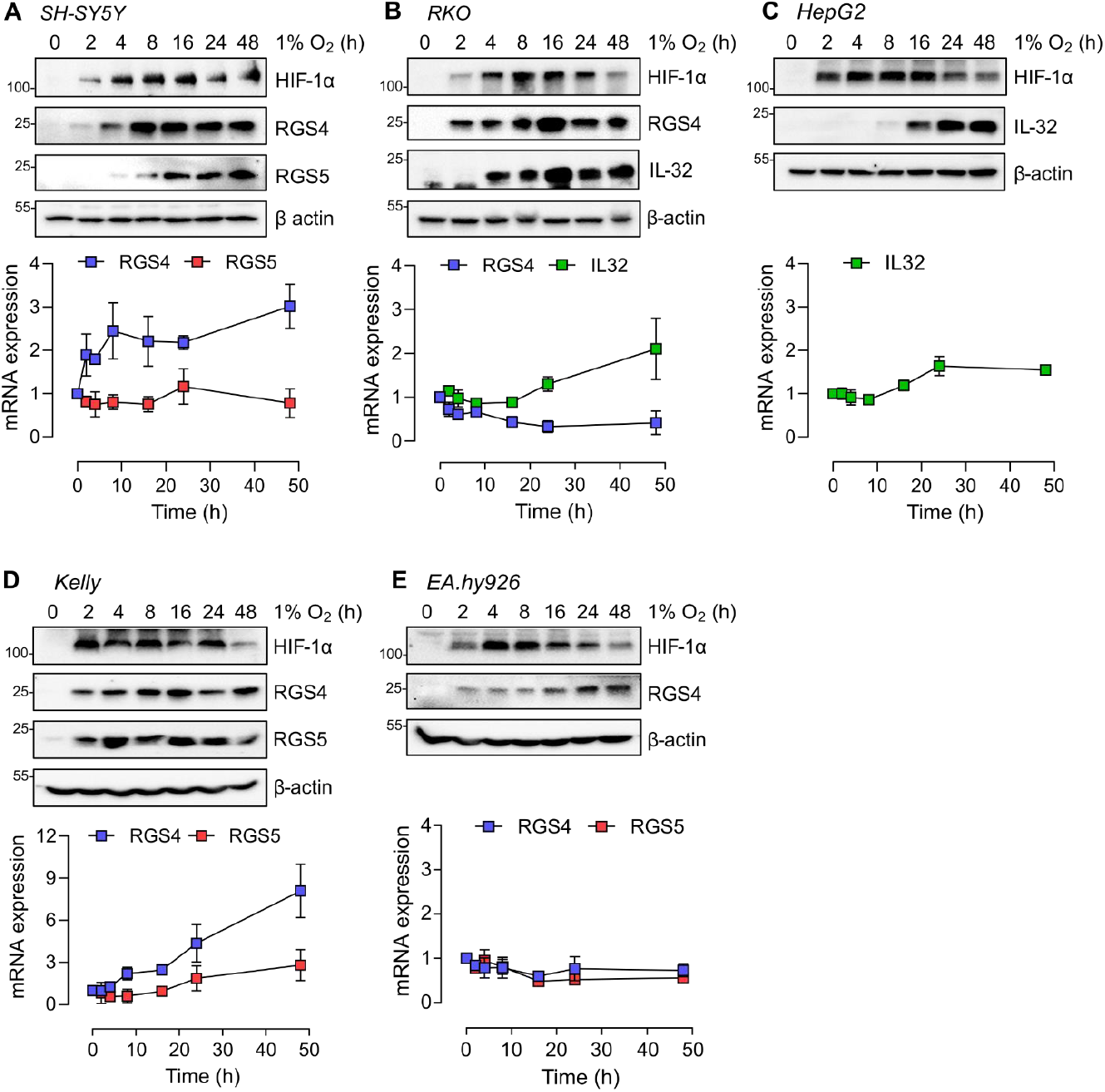
Time course of induction of known hypoxia-inducible proteins. Five cell lines; SH-SY5Y (**A**), RKO (**B**), HepG2 (**C**), Kelly (**D**) and EA.hy926 (**E**) were exposed to 1% O_2_ for the indicated periods of time. Representative immunoblots (*top*) and mRNA levels of the ADO substrates (RGS5, RGS4 and IL-32) are shown below. All data represent the mean ± S.D. from 3 independent experiments.

Since all the ADO targets analysed here have been reported to respond directly or indirectly to the HIF/PHD system in specific settings (30-32), we went on to investigate interactions between these systems. We first examined the expression of *RGS4* mRNA in SH-SY5Y cells in which it is been reported to be a target of HIF as well as ADO (30). siRNA targeting either HIF-1α or -2α was used to confirm the HIF dependency of this response, with successful knock-down confirmed by an attenuation of hypoxia-induced CA9 and VEGFA transcripts, which were selected to report specific HIF-α isoform activity (Fig. 7A). Notably, increased hypoxic induction was observed when cells were treated with siRNA targeting the alternative HIF-α isoform (HIF-2α for CA9 and HIF-1α for VEGFA) which was repressed when both isoforms were targeted. Importantly, hypoxic induction of *RGS4* mRNA was reduced in the presence of siRNA targeting HIF-2α and HIF-1+2α, whilst increased when cells were treated with HIF-1α siRNA alone (Fig. 7A), confirming an action of HIF-2α on *RGS4* transcripts in SH-SY5Y cells. We then proceeded to address the relative contributions of ADO and HIF pathways to the accumulation of RGS4 protein in hypoxia. To this end, ADO-competent or - deficient SH-SY5Y cells were exposed to hypoxia (1% O_2_) in the presence or absence of a HIF-prolyl hydroxylase inhibitor ([(1-chloro-4-hydroxy-isoquinoline-3-carbonyl)-amino]-acetic acid, FG-2216 (33), Fig. 7B), a molecule previously shown not to inhibit ADO (6). As expected, hypoxia resulted in the accumulation of RGS4 in ADO-competent cells. Interestingly however, FG-2216 had no effect on RGS4 protein in normoxic cells whereas in ADO-deficient cells, in which RGS4 was constitutively stable, transcriptional upregulation of mRNA by either hypoxia or FG-2216 resulted in a substantial additional increase in RGS4 protein levels. Thus, combined inhibition of ADO and PHD amplified the induction of RGS4 protein levels. Nevertheless, proteolytic control via ADO likely remains the dominant regulatory mechanism under many conditions, as witnessed by the absence of an effect of FG-2216 in ADO-competent normoxic cells.

**Figure 7.**
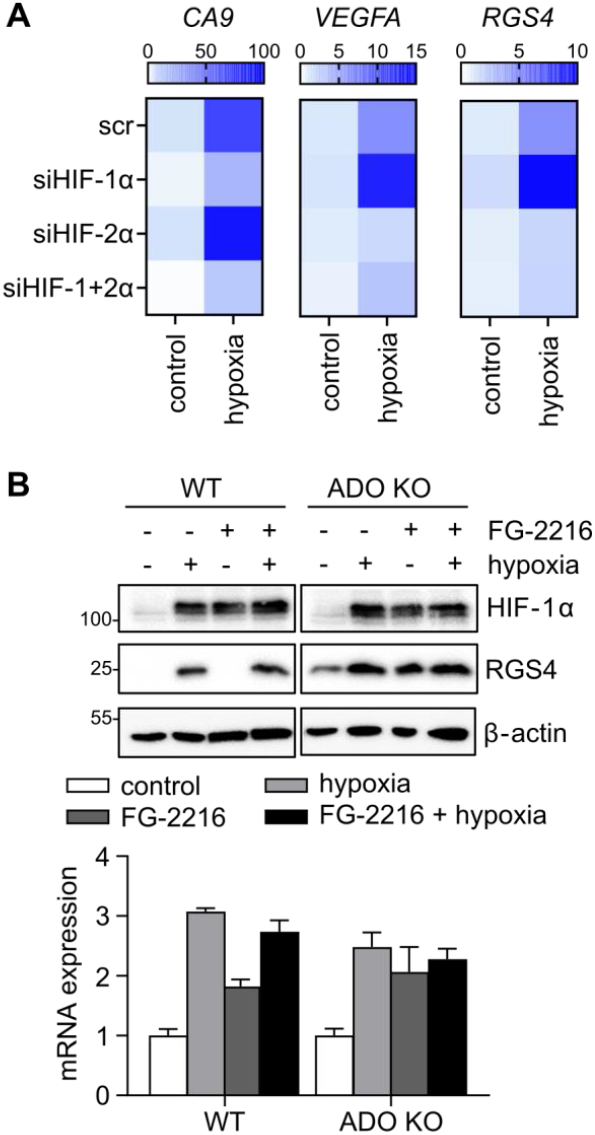
Coordinated regulation of RGS4 by HIF-2α and ADO under hypoxia. (**A**) SH-SY5Y cells were transfected with siRNA targeting HIF-1α and/or HIF-2α, or scrambled control (scr), and subjected to 24h of hypoxia (1% O_2_). RGS4 mRNA levels were assessed alongside canonical HIF-1α (CA9) or HIF-2α (VEGF) target genes as positive controls. The colour scale represents mean fold change relative to cell treated with scr control (normoxia) from 3 independent experiments. (**B**) ADO-competent or -deficient SH-SY5Y cells were exposed to hypoxia (1% O_2_) or treated with the PHD inhibitor FG-2216 (100μM), or both, for 4h and samples blotted for RGS4 and HIF-1α protein. RGS4 mRNA levels were assessed in parallel, confirming transcript upregulation. Immunoblots are representative of 3 separate experiments, and data in the histogram are the mean ± S.D, n=3. All treatments significantly induced RGS4 mRNA, 2-way ANOVA with Holm-Sidak post-hoc analysis, P<0.001.

Conversely, when proteolytic degradation is prevented, accumulation of a given protein will be largely determined by the rate of *de novo* transcription/translation. We therefore wished to explore other interactions between transcriptional regulation and ADO-dependent proteolysis. *IL32* has been shown to be regulated at the transcript level by inflammatory mediators such as TNFα (34,35). We therefore hypothesised that TNFα pre-treatment might influence the rate of IL-32 protein accumulation upon exposure to hypoxia, which was slower in HepG2 cells relative to other substrates in other cell lines (Fig. 6). To test this, HepG2 cells were pre-treated with TNFα for 16 hours to elicit a robust induction of *IL32* mRNA (Fig. 8A), and the effects of timed exposure to hypoxia (1% O_2_, 0.5-8 hours) compared with and without this TNFα pre-treatment. These experiments revealed a marked interaction. Cells pre-treated with TNFα demonstrated a greatly enhanced response of IL-32 protein levels to hypoxia (Fig. 8B) consistent with TNFa amplifying the effects of O_2_-regulated proteolysis transduced by ADO. To confirm this, ADO was inactivated in HepG2 cells by CRISPR/Cas9 gene editing and these cells were treated with TNFα for 16 hours, before a brief exposure to hypoxia (1% O_2_, 2 hours). As ADO has been reported to influence TNFα signalling (36), cells were also treated with IL-1β (37). As shown in Figure 8C, induction of IL-32 protein by TNFα and IL-1β was much higher in ADO-deficient than parental HepG2 cells (Fig. 8C, compare lanes 1-3 and 7-9). As before, induction of IL-32 by hypoxia was greatly exaggerated in cytokine-treated cells, and observed even after brief (2 hours) exposure. However, in ADO-deficient cells the very high levels of IL-32 were not further increased by hypoxia, consistent with ablation of ADO-dependent proteolysis. Taken together these results illustrate strong interactions in the multi-level control of substrates of the ADO pathway, with transcriptional induction amplifying O_2_-dependent regulation of ADO substrates at the protein level.

**Figure 8.**
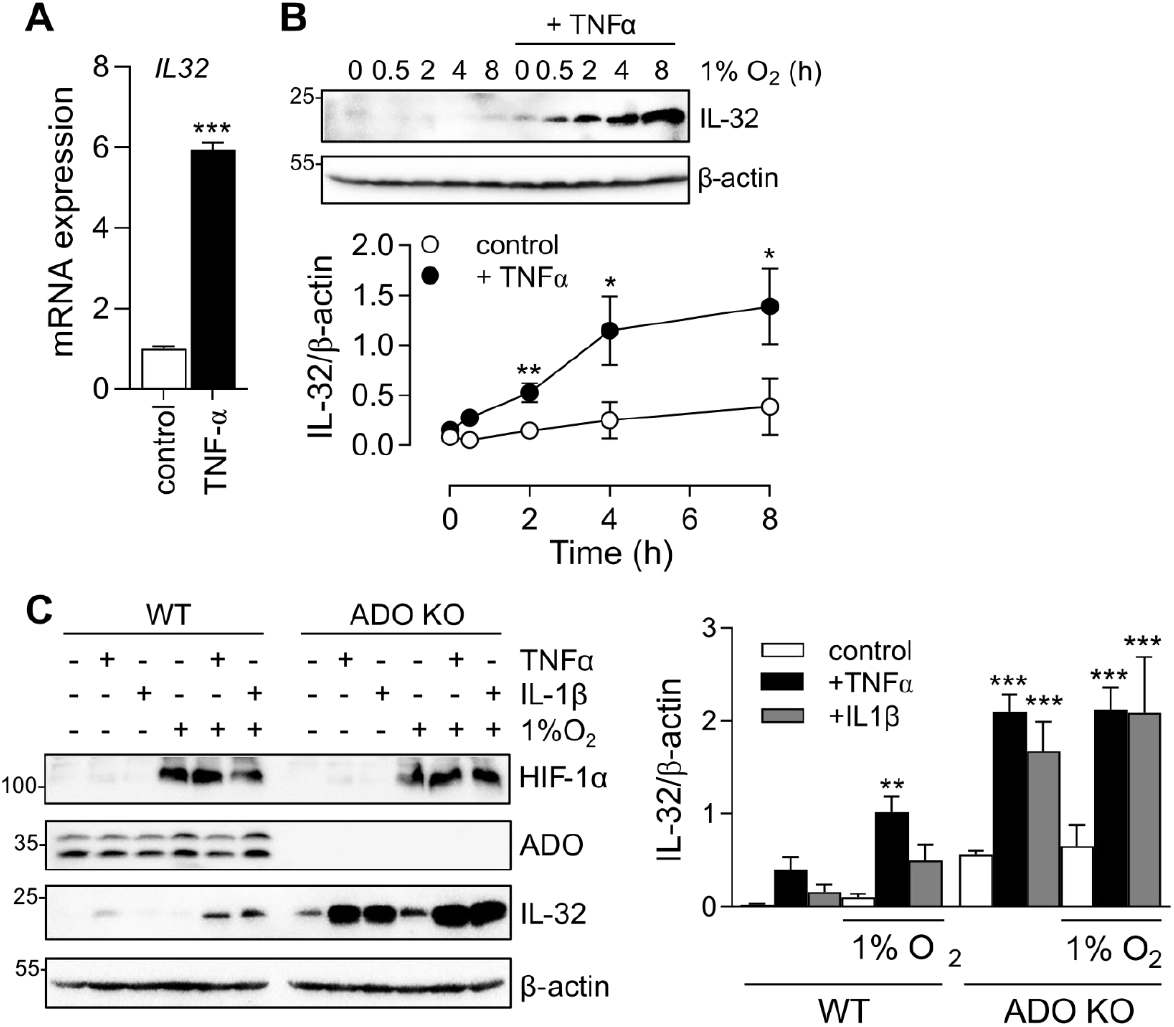
Interplay between transcriptional and proteolytic regulation of IL-32. (**A**) Induction of *IL32* mRNA in HepG2 cells treated for 16h with TNFα (20ng/μL). (**B**) Accumulation of IL-32 protein in HepG2 cells treated with TNFα prior to exposure to hypoxia for the times indicated. (**C**) ADO-competent or -deficient HepG2 cells were treated with TNFα or IL-1β (20ng/μL) for 16h then exposed to hypoxia for a further 2h, and levels of IL-32 protein analysed. HIF-1α protein levels were assayed for comparison. All data represent the mean ± S.D. from 3 independent experiments, *P<0.05, **P<0.01, ***P<0.001, Mann-Whitney T-test (A) or 2-way ANOVA with Holm-Sidak post-hoc analysis (B and C).

## Discussion

Despite a common and fundamental requirement to maintain oxygen homeostasis, many different systems for sensing and responding to hypoxia have been defined in different species (38). Somewhat surprisingly, several of these different systems deploy a common strategy of enzymatic protein oxidation coupled to proteolysis as the core signalling process. Of particular interest are the plant and animal systems that deploy N-terminal cysteine dioxygenation and prolyl hydroxylation to regulate transcriptional responses mediated by ERF-VII transcription factors and HIF respectively. Remarkably, the plant system of N-terminal cysteine dioxygenation is also represented in humans and other animal species by the N-terminal cysteine dioxygenase, ADO, which therefore has the potential to work alongside PHD/HIF. Since oxygen homeostasis must be maintained across different tissues, operating at different levels of oxygen and over different time-scales, we sought to compare and contrast the characteristics of these pathways in human cells. Our findings indicate that the two systems are similarly sensitive to oxygen concentrations, but that their temporal responses and pharmacological interactions display different characteristics. Importantly, direct proteolytic regulation of the ADO targets RGS4 and RSG5 allows for more rapid responses to oxygen, in particular enabling a very rapid offset when cells are re-oxygenated after periods of hypoxia. This may be predicted to be important during rapidly cycling of oxygen levels such as occurs in disorders of breathing control, in ischaemia reperfusion, and in the tumour microenvironment. Similar sensitivity to oxygen also suggested that the PHD/HIF and ADO systems might interact should they share common targets. Remarkably, a number of reports have described transcriptional responses to HIF amongst the limited range of ADO substrates defined to date. Our findings confirmed this and revealed that in specific settings the two systems do indeed interact to amplify responses to hypoxia or inflammatory signals.

These interactions may be important both physiologically and therapeutically where, for instance, HIF prolyl hydroxylase inhibition may be more effective in hypoxic regions where ADO activity is also reduced.

Although both PHD and ADO enzymes share similar kinetic properties relating to O_2_, the reactions they catalyse are quite distinct. HIF prolyl-hydroxylation requires 2-oxoglutarate as co-substrate and hence are potentially affected by metabolic signals that modulate cellular levels of this metabolite. The N-terminal cysteine dioxygenase ADO incorporates both oxygen atoms into the cysteine residue and has no such co-substrate requirement or reaction by-products. However, in common with the PHD enzymes, ADO deploys a His-Asp-His facial triad to co-ordinate the catalytic Fe^2+^, which potentially exchanges with cellular iron and other metals. Although our studies revealed that ADO, like the PHDs, is sensitive to inhibition by iron chelating agents, these effects, and those of transition metals ions, differed substantially between the two types of enzyme. Notably, whereas inhibition of the PHDs by chelating agents was broadly concordant with their action to reduce intracellular iron, this was not the case with inhibition of ADO, for which only selected chelators were effective and inhibitory activity did not correlate with simple depletion of intracellular iron. This implies that the catalytic Fe^2+^ in ADO is not freely exchanging with cytosolic Fe^2+^ and that the inhibitory chelators have a more specific interaction with the catalytic iron centre of ADO, or, conceivably, that their inhibitory activity is unrelated to the chelation of iron. Of note, all chelators demonstrated here to inhibit ADO activity *in cellulo* (2,2DIP, Dp44mT and SIH) have been described as selective towards Fe^2+^, whereas DFO, ICL670A and L1 all appear to preferentially bind Fe^3+^ (39,40).

In keeping with restricted exchange of Fe^2+^ into ADO, we obtained little evidence that this enzyme was sensitive to transition metal ions. Interestingly however, although only minimal induction of an ADO-dependent reporter gene was observed, Co^2+^ ions induced striking upregulation of RGS4 and RGS5 proteins, as has recently been reported by others (25). It was proposed that Co^2+^-induced trioxidation of the N-terminal cysteine of RGS4/5 leads to lysosomal degradation rather than via the canonical 26S proteasome, with the former presumably being much less efficient to account for the observed large increase in accumulated RGS4/5 in cells exposed to Co^2+^. If true, a reduction in the steady-state RGS4/5 protein level in ADO-deficient cells treated with Co^2+^ would be expected, as a portion of the constitutively stabilised RGS4/5 would now be targeted for autophagy and a new equilibrium achieved. We have presented several lines of evidence in contradiction to this hypothesis (Fig. 4). The near absence of an action of Co^2+^ on the RGS4_1-11_GFP reporter protein and the persistence of an increase in endogenous RGS4/5 in ADO-deficient cells and on a C2A mutant of RGS4 all suggest that the action of Co^2+^ on RGS4 and RGS5 are not, in the main, mediated by an effect on the oxidation status of N-terminal cysteine. We suggest that the action of N-acetyl cysteine to inhibit this proposed oxidation likely results from direct chelation of Co^2+^ ions in solution (41,42).

Indeed, NAC is clinically approved for the treatment of acute Co^2+^ poisoning (43,44). Further work is needed to define the precise mechanism through which Co^2+^ acts to increase RGS4/5 stabilisation, and its relevance to the toxicology of cobalt.

In summary our data provide evidence that ADO acts as a physiological oxygen sensor operating in cells alongside, and interacting with, HIF. The ADO system responds at oxygen concentrations similar to those that induce HIF, but its direct action to promote degradation of RGS proteins over a time-scale of minutes enables the transduction of responses to altered oxygen levels on a shorter time-scale. *In cellulo* characterisation of the ADO pathway revealed marked interactions with the transcriptional induction of specific targets and distinct divalent metal pharmacology that offers new insight into the biochemistry of cellular dioxygenases.

### Experimental procedures

#### Cell culture

The human cell lines SH-SY5Y, RKO and EA.hy926 were cultured as described previously (6). HepG2 were kindly gifted by Jane McKeating (University of Oxford, UK) and Kelly cells were purchased from the European Collection of Authenticated Cell Cultures (ECACC 92110411). RKO, EA.hy926 and HepG2 cells were cultured in DMEM, and SH-SY5Y and Kelly cells in DMEM/F12, all supplemented with 10% fetal bovine serum, 2mM L-Glutamine and 100 U/ml penicillin/10μg/ml streptomycin. All cell lines were maintained at 37°C incubator containing 5% CO_2_, and hypoxic exposure performed using an atmosphere-regulated workstation set to 0.1-7.5% O_2_: 5% CO_2_: balance N_2_ (Invivo 400, Baker-Ruskinn Technologies).

### Immunoblotting

Protein samples were collected in lysis buffer (10 mM Tris pH 7.5, 0.25 M NaCl, 0.5% Igepal) supplemented with Complete™ protease inhibitor cocktail (Sigma Aldrich) and centrifuged at 13,000rpm for 3 minutes at 4°C. The supernatant was mixed with Laemmli sample buffer and proteins separated via SDS-PAGE electrophoresis. Membranes were blocked in 4% milk for 1 hour, then incubated in primary antibody overnight: HIF-1α (610959, BD Biosciences), RGS5 (sc-514184, SCBT), CA9 (5648, CST), RGS4 (15129, CST), IL-32 (sc-517408, SCBT), ADO (ab134102, Abcam), ATE1 (HPA038444, Human Protein Atlas), PHD3 (188e (45)) GFP (11814460001, Sigma Aldrich), and HA (3F10, Roche). HRP-conjugated secondary antibodies were sourced from DAKO and used in conjunction with chemiluminescence substrate (West Dura, 34076, Thermo Fisher Scientific) to visualise protein expression using a ChemiDoc XRS+ imaging system (BioRad). β-actin primary antibody was conjugated directly to HRP (ab49900, Abcam). Densitometric analysis was performed using ImageJ software (NIH) and values presented relative to β-actin.

### RT-qPCR

mRNA was extracted from Tri-Reagent lysates by phase separation and equal amounts were used for cDNA synthesis using the High-Capacity cDNA Kit (Applied Biosystems). qPCR analysis was performed using Fast SYBR Green Master Mix on a StepOne thermocycler (Thermo Fisher Scientific) using the ΔΔCt method. Levels of the housekeeping gene Hypoxanthine-guanine phosphoribosyl transferase (HPRT) were used as a reference. Sequences for the primers used are as follows;

RGS4 (F_*GCAAAGGGCTTGCAGGTCT, R_CAGCAGGAAACCTAGCCGAT)*,

*RGS5(F_TGGTGACCTTGTCATTCCG, R_TTGTTCTGCAGGAGTTTGT)*,

*IL32 (F_CTTCCCGAAGGTCCTCTCTGAT, R_GTCCTCAGTGTCACACGCT)*,

HPRT(F_*GACCAGTCAACAGGGGACAT, R_AACACTTCGTGGGGTCCTTTTC)*,

*ADO(F_GCCGGGACTGCCACTATTAC, R_ACCAGAAGTCATCGGCCTGT)‘*,

*ATE1(F_CTGATTTGCTGTGCCCTGAG, R_GGTTCCGTACTGCGATCCTC*,

*EGLN1(F_GCAGCATGGACGACCTGATA, R_CCATTGCCCGGATAACAAGC)*,

*EGLN3 (F_CACGAAGTGCAGCCCTCTTA), R_TTGGCTTCTGCCCTTTCTTCA)*,

*METAP1(F_CATCAAGCTGGGCATCCAGG, R_GCTTCGCCTTTTCATCTTTTGC)*,

*UBR1(F_TATGGAGGAAGAGAGCACCCC, R_GATGGACCCCGTTTAGGACC*)

*UBR2(F_ACCAGCAGTTGCAGAGAGAT, R_GTGATGAGCATTCGAGCCAGA)*,

*CA9(F_CTTGGAAGAAATCGCTGAGG, R_TGGAAGTAGCGGCTGAAGTC)*

*VEGFA (F_TGTCTAATGCCCTGGAGCCT, R_GCTTGTCACATCTGCAAGTACG)*

### siRNA-mediated gene knockdown

SH-SY5Y cells were seeded into 12-well plates and transfected twice on subsequent days with 40nM of either scrambled siRNA (4390843) or siRNA sequences targeting HIF-1α (s6539), HIF-2α (s4700, Ambion) or a combination of both, diluted in OptiMEM and Lipofectamine RNAiMAX (Thermo Fisher Scientific, UK).

### Monitoring ADO activity using RGS4_1-11_GFP

The design of the ADO reporter construct was based on a derivative previously published (RGS4_1-20_GFP) (6,20). The first 11 amino acids of human RGS4 (Uniprot: P49798-1) were fused to the N-terminus of eGFP. This construct was then inserted into the pcDNA3+ vector and 2μg of plasmid DNA transfected into a 6cm plate of subconfluent RKO cells using GeneJuice transfection reagent. Transfected cells were selected for using 1.4mg/ml G418 for 2 weeks, producing a polyclonal pool of reporter cells. Cells were seeded at 10,000 cells/well in black 96-well plates and allowed to grow to confluence, before being treated with increasing concentrations of Fe^2+^ chelator (2,2DIP -2,2′-dipyridyl, DFO – deferoxamine, Dp44mT - di-2-pyridylketone 4,4-dimethyl-3-thiosemicarbazone, ICL670A - desferasirox, L1 – deferiprone, SIH - salicylaldehyde isonicotinoyl hydrazone) or divalent metal ion for 24 hours, after which time fluorescence was read at 480nm_ex_/520nm_em_ using a plate reader (FLUOstar, BMG Labtech). Cells were then incubated with MTT (0.5mg.ml^-1^) for 4 hours to monitor viability (46).

### Measurement of Intracellular and ADO/PHD2-bound Fe^2+^

Intracellular labile Fe^2+^ levels were measured using FerroOrange, a fluorescent probe specific to Fe^2+^ (47). RKO cells were seeded in black 96-well plates and treated with Fe^2+^ chelators for 4 hours in standard medium, which was subsequently replaced with imaging buffer (in mM; 117 NaCl, 4.5 KCl, 25 NaHCO_3_, 11 Glucose, 1 MgCl_2_, 1 NaH_2_PO_4_, 1 CaCl_2_) containing 1μM FerroOrange. Cells were incubated in probe containing medium for 30 minutes at 37°C then fluorescence measured at 544nm_ex_/590nm_em_ using a FLUOstar Omega plate reader (BMG Labtech, UK). Fluorescence from a well containing medium alone (no FerroOrange) was used as a blank. To measure total iron content of ADO and PHD2, N-terminally FLAG-tagged enzymes were immunoprecipitated from cells treated with either 2,2-dipyridyl or DFO (both 100μM) and total Fe content analysed by Inductively coupled plasma mass spectrometry (ICP-MS). SH-SY5Y cells stably expressing an N-terminally FLAG-tagged *Hs*ADO or *Hs*PHD2 were cultured to confluence in 15cm plates, then treated for 4 hours with 100μM of either DFO or 2,2DIP and lysed in lysis buffer (see above). Enzymes were immunoprecipitated using anti-FLAG(M2) conjugated agarose beads overnight at 4°C, washed in PBS five times then eluted in 2% nitric acid. Beads incubated with lysis buffer (without cellular extract) were subjected to the same procedure as background readings for correction. Elutions were further diluted (2X) in 2% nitric acid and analysed on a Perkin Elmer NexION 2000B ICP-MS, calibrated using external calibration analysis with QMX standard dilutions and spiked with 1ng/g Rhodium to normalise any instrument drift. Values in immunoprecipitates were corrected for non-specific background by subtracting the iron detected in equivalent samples from cells that did not express any FLAG-tagged construct.

### Cobalt-pull down

To measure the Co^2+^ binding of RGS4/5, ADO-deficient SH-SY5Y cell lysates were incubated with HisBind resin (Millipore 69670) pre-charged with Co^2+^ ions, allowing the precipitation of Co^2+^-binding proteins as described previously (28). 25μL (packed volume) of HisBind resin was washed in PBS and then incubated with 50mM CoCl_2_ at room temperature for 1 hour, then washed 5 times with lysis buffer. Cell lysate (15cm dish per condition) was then incubated with Co^2+^ (or control) loaded resin for 1 hour under constant rotation at 4°C. Following this, resin was pelleted (2000 rpm, 2 minutes) and a sample taken and mixed with sample buffer to represent an unbound fraction. The resin was then further washed and subjected to the following elutions; 60μL 60mM imidazole for 10 minutes, 60μL 1M imidazole, then finally 70μL SDS sample buffer. After each elution, the supernatant was taken and heated at 95°C for 5 minutes, then subjected to SDS-PAGE and immunoblotted as described above.

## Data Availability

All data are contained within the manuscript

